# Prediction of SARS-CoV interaction with host proteins during lung aging reveals a potential role for TRIB3 in COVID-19

**DOI:** 10.1101/2020.04.07.030767

**Authors:** Diogo de Moraes, Brunno Vivone Buquete Paiva, Sarah Santiloni Cury, João Pessoa Araújo Junior, Marcelo Alves da Silva Mori, Robson Francisco Carvalho

**Affiliations:** Department of Structural and Functional Biology, Institute of Biosciences, São Paulo State University (UNESP), Botucatu, São Paulo, Brazil; Faculty of Medicine, São Paulo State University, UNESP, Botucatu, São Paulo, Brazil; Department of Chemical and Biological and Sciences, Institute of Biosciences, São Paulo State University (UNESP), Botucatu, São Paulo, Brazil; Department of Biochemistry and Tissue Biology, Institute of Biology, State University of Campinas (UNICAMP), Campinas, SP, Brazil

**Author notes:** These authors contributed equally to this work (Diogo de Moraes, Brunno Vivone Buquete Paiva). **Corresponding author information** Robson Francisco Carvalho, Department of Structural and Functional Biology, Institute of Biosciences, São Paulo State University (UNESP), CEP: 18.618-689, Botucatu, São Paulo – Brazil.

**Keywords:** COVID-19, SARS-CoV-2, tribbles homolog 3, α-hydroxylinoleic acid, lung aging

## Abstract

COVID-19 is prevalent in the elderly. Old individuals are more likely to develop pneumonia and respiratory failure due to alveolar damage, suggesting that lung senescence may increase the susceptibility to SARS-CoV-2 infection and replication. Considering that human coronavirus (HCoVs; SARS-CoV-2 and SARS-CoV) require host cellular factors for infection and replication, we analyzed Genotype-Tissue Expression (GTEx) data to test whether lung aging is associated with transcriptional changes in human protein-coding genes that potentially interact with these viruses. We found decreased expression of the gene tribbles homolog 3 (*TRIB3*) during aging in male individuals, and its protein was predicted to interact with HCoVs nucleocapsid protein and RNA-dependent RNA polymerase. Using publicly available lung single-cell data, we found *TRIB3* expressed mainly in alveolar epithelial cells that express SARS-CoV-2 receptor ACE2. Functional enrichment analysis of age-related genes, in common with SARS-CoV-induced perturbations, revealed genes associated with the mitotic cell cycle and surfactant metabolism. Given that TRIB3 was previously reported to decrease virus infection and replication, the decreased expression of *TRIB3* in aged lungs may help explain why older male patients are related to more severe cases of the COVID-19. Thus, drugs that stimulate TRIB3 expression should be evaluated as a potential therapy for the disease.

## Introduction

The first cases of infections with the severe acute respiratory syndrome coronavirus 2 (SARS-CoV-2) in humans were identified in December 2019 in Wuhan, China^1,2^, and since then, the coronavirus disease 2019 (COVID-19) rapidly became pandemic^3^. Studies have shown that older individuals with comorbidities are associated with more severe cases of COVID-19^4^. These patients are more likely to develop pneumonia and respiratory failure due to alveolar damage^5,6^, suggesting that lung aging impacts disease progression and mortality.

SARS-CoV-2 requires host cellular factors for successful infection and replication^7^. For example, angiotensin-converting enzyme 2 (ACE2) is the receptor for the SARS-CoV-2 spike protein receptor-binding domain (RBD) for viral attachment^7,8^. The conserved evolutionary relationship between the 2019 novel SARS-CoV-2 and SARS-CoV^9^ opens up the possibility to explore the relationships between these human coronaviruses (HCoVs) in public databases. Computational predictions of SARS-CoV–human protein-protein interactions (PPIs) may identify mechanisms of viral infection and drug targets^9–11^. In this context, the Genotype-Tissue Expression (GTEx) database^12,13^ has provided insights into age-related genes^14,15^ and, associated with single-cell transcriptomics, could predict SARS-CoV-2–PPIs in aging lungs. This characterization is crucial for older adults, which are more vulnerable to the disease^16–18^. Thus, we analyzed whether lung aging is associated with transcriptional changes in proteins that potentially interact with SARS-CoV-2.

We first identified differentially expressed genes (DEGs) during aging in GTEx human lung samples (release V7) (Data S1). The numbers of significant DEGs increased with aging (Log fold-change ≥ |1| and FDR < 0.05), and individuals of 60-69 year old (yo) presented the highest number of DEGs, in comparison to young adults (20-20 yo) (Figures S1-S2, Table S1). Clustering of these DEGs identified age-associated profiles (Figure 1a). Among the transcripts translated into proteins predicted as interacting with SARS-CoV, the hyaluronan and proteoglycan link protein 2 (*HAPLN2*) increased with aging, while tribbles homolog 3 (*TRIB3*) decreased (Figure 1b, Table S2). HAPLN2 was predicted to interact with virus proteins spike glycoprotein and E2 glycoprotein precursors, while TRIB3 with nucleocapsid protein and RNA-dependent RNA polymerase (Figure 1b; Table S3). Importantly, the SARS-CoV-2 nucleocapsid protein has a sequence identity of 89.6% compared to SARS-CoV^9^. The expression of *TRIB3* also decreased in the lung, specifically in males with ≥ 40 years (Figure 1c), in a cohort with additional samples (GTEx, release V8).

**Figure 1.**
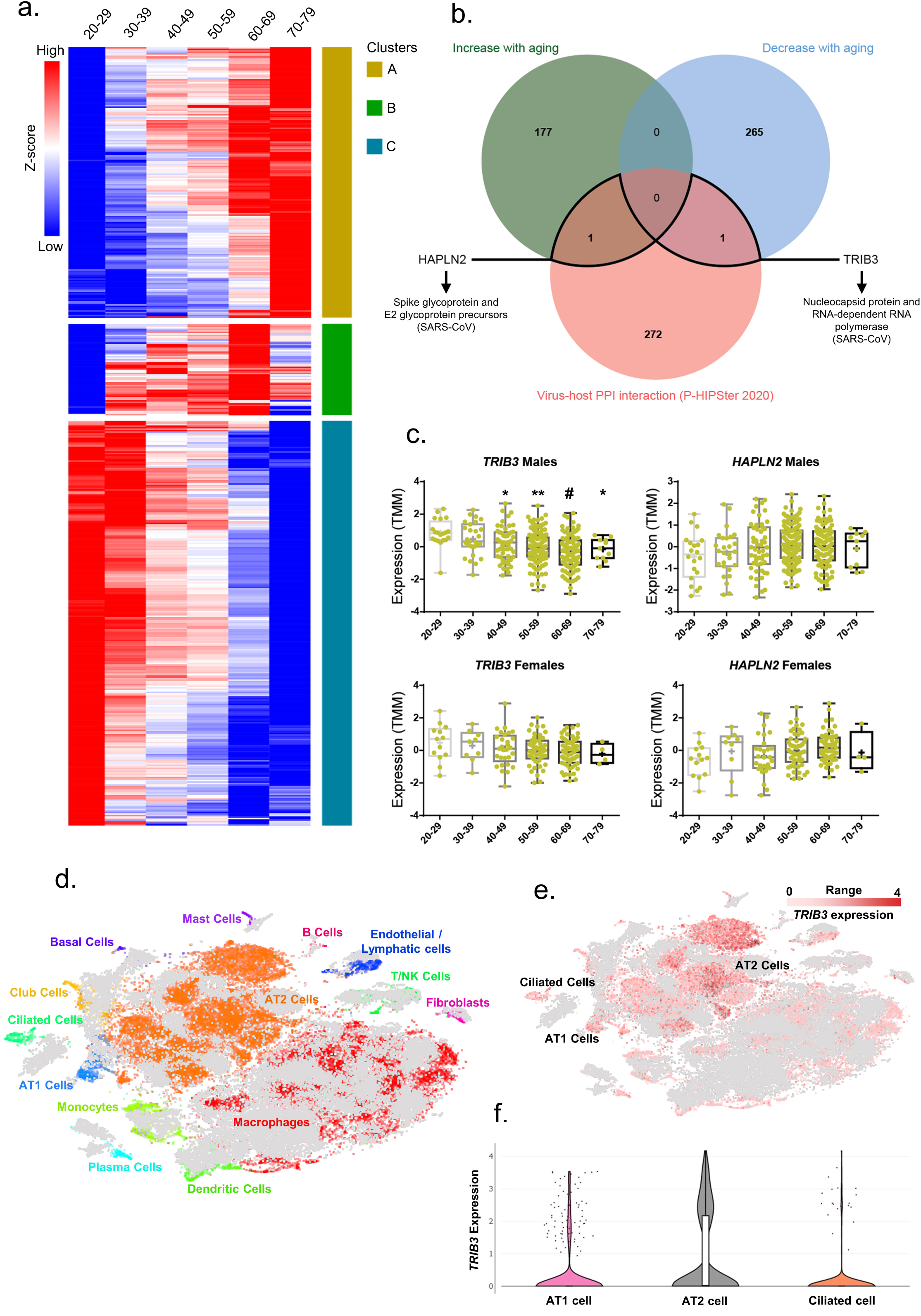
Lung gene expression of TRIB3, a protein that potentially interacts with SARS-CoV-2 proteins, decreases in male individuals during aging. a) Heatmap of genes found as differentially expressed (DEGs; mean expression) in, at least, one age-group when compared to young adults (20-29 yo). Rows were clustered using Euclidian distance. Clusters A and C contain genes that increase or decrease with age, respectively. b) Venn diagram of DEGs during aging shared with the corresponding proteins that potentially interact with SARS-CoV-2 (arrows) c) Boxplot of gene expression levels (TMM). * P < 0.05, ** P < 0.001, and # P < 0.0001: statistical significance vs. young adults (20-29 yo). d) Unsupervised clustering of human lung cell populations identified in non-diseased lung samples in the *t*-distributed Stochastic Neighbor Embedding (tSNE) plot. Grey dots: removed cells (disease). e) *TRIB3* expression in lung cell populations. f) Violin plots of *TRIB3* expression levels in lung single-cells.

The reanalysis of lung single-cell RNA sequencing data ^19,20^ demonstrated *TRIB3* expressed mainly in alveolar type I (AT1) and type II (AT2) cells and in ciliated cells (Figure 1d-f, Figure S3), which also express SARS-CoV-2 receptor ACE2^7,8,21^. The involvement of TRIB3 in viral infection is poorly understood; however, its inhibition was associated with an increase of hepatitis C virus (HCV) replication^22^. Additionally, TRIB3 negatively regulates the entry step of the HCV life cycle and propagation^22^ and thus may constitute a common protective host factor for other positive-sense single-strand RNA viruses. *TRIB3* is also one of the unfolded protein response (UPR)-related genes with the strongest positive correlation with the intracellular abundance of the flavivirus dengue and Zika^23^. Considering the need for drugs to treat COVID-19, the α-hydroxylinoleic acid (ABTL0812) induces the expression of TRIB3 by inhibiting the PI3K/AKT/mTOR axis and promoting autophagy cell death in cancer^24^. We highlight that the lifecycle of coronaviruses depends on several host-cell encoded cellular pathways, and among these pathways, UPR and autophagy pathways of the host cells are essential to the life cycle of coronaviruses^25^.

Finally, we compared SARS-CoV-induced perturbations in host gene expression, from public Gene Expression Omnibus (GEO) datasets, with our list of DEGs in GTEx lung samples during aging (Figure 2, Table S8). We found that genes that decrease their expression with aging and genes that are up-regulated with SARS-CoV infections generated the most significant network, with over-represented genes associated with mitotic cell cycle and surfactant metabolism (Figure 2). The decreased capacity of cellular division on aging is associated with cellular senescence - a mechanism that stops cells with damaged DNA from replicating^26^ - and progenitor cell exhaustion^27^. The altered metabolism or secretion of surfactants by AT2 cells reduces the ability of the lungs to expand and increases the risk of lung collapse in HCoVs infections^28,29^. Moreover, *Sftpc*^-/-^ (Surfactant Protein C) mice have worse viral infections than controls^30^, and its human homolog decreased with aging while it is up-regulated on SARS-CoV infections (Figure 2). Thus, the pneumonia-like lung injury found on severe cases of COVID-19 infections^5,6^ may be aggravated by impaired lung regeneration and altered metabolism of surfactants in older male patients.

**Figure 2.**
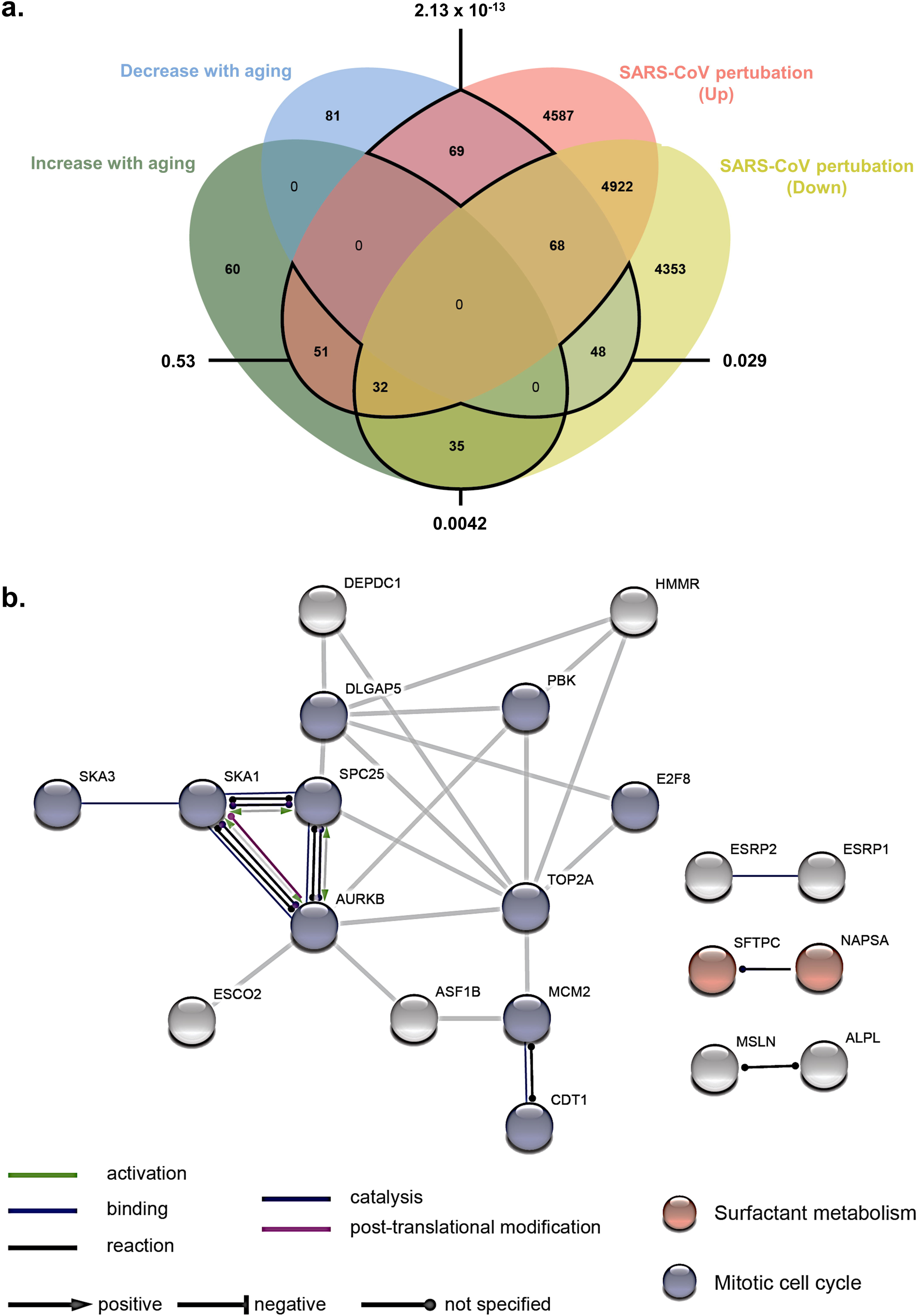
a) Venn diagram of differentially expressed genes (corresponding proteins) during aging shared with SARS-CoV-induced perturbations in host gene expression. Values outside the diagram: PPI enrichment P-value. b) PPI network based on the genes that decreased expression during aging and are up-regulated in SARS-CoV-induced perturbations in host gene expression. Table S8 contains the complete list of over-represented terms.

Although the genes and pathways we highlighted here were identified based on a robust statistical significance, we emphasize that other methods of over-time gene expression analyses applying different cutoffs could be considered; using GTEx V8 cohort or separating males and females may result in different sets of age-related genes in the lung. Further analyses should be conducted to identify more functional differences between male and female lungs during aging. Moreover, clinical data from these individuals - such as diabetes or heart disease - important factors influencing COVID-19 outcome - were not evaluated. However, due to the nature of GTEx donor consent, public phenotypes (including clinical data) are limited. Access to the protected phenotypes of the individuals needs an application via dbGaP (Genotypes and Phenotypes database), which, associated with reanalysis of the transcriptomics data, may take a significant amount of time. Part of the results presented herein derives from a previously unpublished paper focusing on aging lung on a different topic. Nevertheless, we decided to release this data focusing on SARS-CoV-2, due to the emergency of the current pandemic.

In conclusion, we show that lung gene expression of TRIB3, a protein predicted to interact with the nucleocapsid protein and the RNA-dependent RNA polymerase of HCoVs, decreases specifically in males during aging. This study provides insights into aging and COVID-19 based on the transcriptional profile of the aging lung and reveals a potential role for TRIB3, surfactant metabolism, and mitotic cell cycle. Considering that TRIB3 may decrease virus infection and replication, strategies to stimulate TRIB3 expression should be tested for the treatment of COVID-19.

## Declaration of Interest

The author declares no conflicts of interest.

## Ethical Approval

Not applicable as we used publicly available data.

## Acknowledgments

This research was supported by the National Council for Scientific and Technological Development, CNPq (Process 311530/2019-2 to RFC, and scholarship #870415/1997-2 to SSC), by the Coordenação de Aperfeiçoamento de Pessoal de Nível Superior, CAPES, Brasil, Finance Code 001 (scholarship to DM), and by Fundação de Amparo à Pesquisa do Estado de São Paulo (2017/01184-9 to MAM). The results shown here are, in part, based upon data generated by the Genotype-Tissue Expression project (GTEx) (https://gtexportal.org/).

## Methods

### Genes differentially expressed in GTEx lung samples during aging

The RNA-Seq data analysis of lung tissues was performed using 427 samples from males and females available at the GTEx portal (release V7) (https://www.gtexportal.org/)^31^. We first used the BioJupies platform (https://amp.pharm.mssm.edu/biojupies/)^32^ to identify the differentially expressed genes (DEG) in lung samples during aging. The expression data of each subject was distributed according to their age range: 30-39; 40-49; 50-59; 60-69 and 70-79 years old (yo), and each age range was compared with the group of young adults (20-29 yo) as a common control (Data S1, Supplementary Table S9. Genes with Log2 of fold change ≥ |1| and false discovery rate (FDR) < 0.05 were considered as differentially expressed (DEGs).

Next, we performed a hierarchical clustering analysis based on Pearson correlation for age ranges using the mean expression (Trimmed Mean of M-values, TMM) of the DEGs found previously in, at least, one age range. This analysis aimed at the identification of clusters with gradients of gene expression across age ranges, showing increased or decreased expression during aging (method adapted from Theunissen et al., 2011^33^). The clusterization of gene expression profiles on female lung samples generated clusters with less evident gradients compared to the males (Figure 1a and Figure S4). Thus, considering the clear clusterization found in males, and the fact that older males seem to have a worse prognosis on COVID-19^6^, we focused on male profiles to identify age-related lung genes.

Our research group previously used the strategy described above to analyze RNA-Seq data of GTEx lung samples (release V7) during aging. We reutilized these results due to the urgency of the current pandemic situation (see limitations section). With the availability of the last release of the GTEx dataset (release V8), which presents a higher number of lung samples with transcriptomic data from individuals of both sexes (Table S10), we used this last available cohort (V8) for further gene-specific analyses. The expression of *HAPLN2* and *TRIB3* (TMM normalized; V8 cohort) were compared using One-way ANOVA followed by Dunnett’s multiple comparisons test, with the GraphPad Prism version 6.00 for Windows (GraphPad Software, La Jolla, California, USA). Differences with a P-value < 0.05 were considered significant.

#### Predicted virus-host protein-protein interactions based on lung genes that increase or decrease expression during aging and SARS-CoV-induced perturbations

The conserved evolutionary relationship between the 2019 novel SARS-CoV-2 and SARS-CoV^9^ opens up the possibility to explore the relationships of these human coronaviruses (HCoVs) in publicly available databases. Thus, we compared the list of DEG in the lung during aging with corresponding human proteins that potentially interact with HCoVs (Table S2). The HCoVs-human PPIs were obtained using data from the Pathogen–Host Interactome Prediction using Structure Similarity (P-HIPSTer, http://phipster.org/) database, which is a comprehensive catalog of the virus– human PPIs predicted based on protein structural information^11^ (Table S2). P-HIPSTer has an experimental validation rate of ∼76%^11^. We also compared the list of DEG with recently added libraries for virus perturbations (up- and down-regulation) from GEO datasets (GSE33266, GSE50000, GSE49262, GSE50878, GSE49263, GSE40824, GSE50878, GSE49263, GSE47960, GSE47961, GSE47962, GSE17400, and GSE40824), available at the EnrichR database^34^. Access in March 2020. Genes that are common in either direction of both conditions were analyzed on STRING (https://string-db.org/) ^35^. Access in March 2020.

#### Single-cell analysis of human lung datasets

We analyzed the expression of *TRIB3, HAPLN2*, and *ACE2*, in different lung cell populations by using two previously published human single-cell RNA-seq data (Table S4)^19,20^. The first dataset^20^ was explored in the UCSC Cell Browser (http://nupulmonary.org/resources/), aiming the identification of the cell populations expressing those genes. The samples with pulmonary fibrosis presented in this dataset were omitted from our analysis, and only non-diseased lung samples were included (n=8). We further used an independent single-cell RNA-seq dataset^19^ (n=5), available at the Human Cell Atlas Portal (https://data.humancellatlas.org/explore/projects/c4077b3c-5c98-4d26-a614-246d12c2e5d7), to confirm that expression *TRIB3* and *ACE2* are expressed in alveolar epithelial cells (types 1 and 2) and in ciliate cells. Access in March 2020.

#### Protein-Protein Interactions (PPI) networks based on lung genes that increase or decrease expression during aging

The corresponding proteins of the DEG shared with the list of DEG from libraries for virus perturbations were queried in the STRING database (Search Tool for Retrieval of Interacting Genes, version 10.5; https://string-db.org/)^35^, for the construction of PPI networks. We considered the following settings: text mining, experiments, databases, and co-expression as sources of active interaction. We selected the minimum interaction score of 0.900 (highest confidence), and the disconnected nodes were hidden to simplify the display. We evaluated the PPI enrichment P-values, which verifies the number of interactions of a set of proteins compared with a random set of similar size. The PPI enrichment P-value represents the statistical significance provided by STRING. Access in March 2020.

#### Data representation and analysis

The clustering analyses of the expression profiles were performed using the web tool Morpheus (https://software.broadinstitute.org/morpheus)^36^. Venn diagrams were plotted using the Jvenn online tool (https://jvenn.toulouse.inra.fr)^37^. Volcano Plots were constructed using the web tool: https://paolo.shinyapps.io/ShinyVolcanoPlot/.

## Data Availability Statement

All data is available in the manuscript.

## List of Supplementary Tables and Supplementary Figures

**Supplementary Data 1**

**Supplementary Table 1**. Differentially expressed genes in lung tissue from male individuals during aging, compared to 20-29 yo individuals (Log2FC ≥ |1|; FDR <0.05) and the clusters of genes that increased or decreased with aging.

**Supplementary Table 2**. Human-SARS-CoV Interactome based on the in silico computational framework P-HIPSTer (http://phipster.org/).

**Supplementary Table 3**. The Human-SARS-CoV interactome was obtained in the P-HIPSTer (http://phipster.org/) for aging lungs.

**Supplementary Table 4**. Characterization of the lung transplant donors used for single-cell transcriptome analysis.

**Supplementary Table 5**. Datasets for SARS-CoV from the library Virus Perturbations from GEO-up (EnrichR; http://amp.pharm.mssm.edu/Enrichr/).

**Supplementary Table 6**. Datasets for SARS-CoV from the library Virus Perturbations from GEO-down (EnrichR; http://amp.pharm.mssm.edu/Enrichr/).

**Supplementary Table 7**. Up- and down-regulated genes for SARS-CoV from the libraries for virus perturbations from GEO (EnrichR; http://amp.pharm.mssm.edu/Enrichr/).

**Supplementary Table 8**. Gene ontology enrichment analysis based on the STRING database.

**Supplementary Table 9**. The number of GTEx lung samples (release V7) used for gene expression analysis according to age group.

**Supplementary Table 10**. The number of GTEx lung samples (release V8) used for gene expression analysis according, age group and gender.

**Figure S1.**
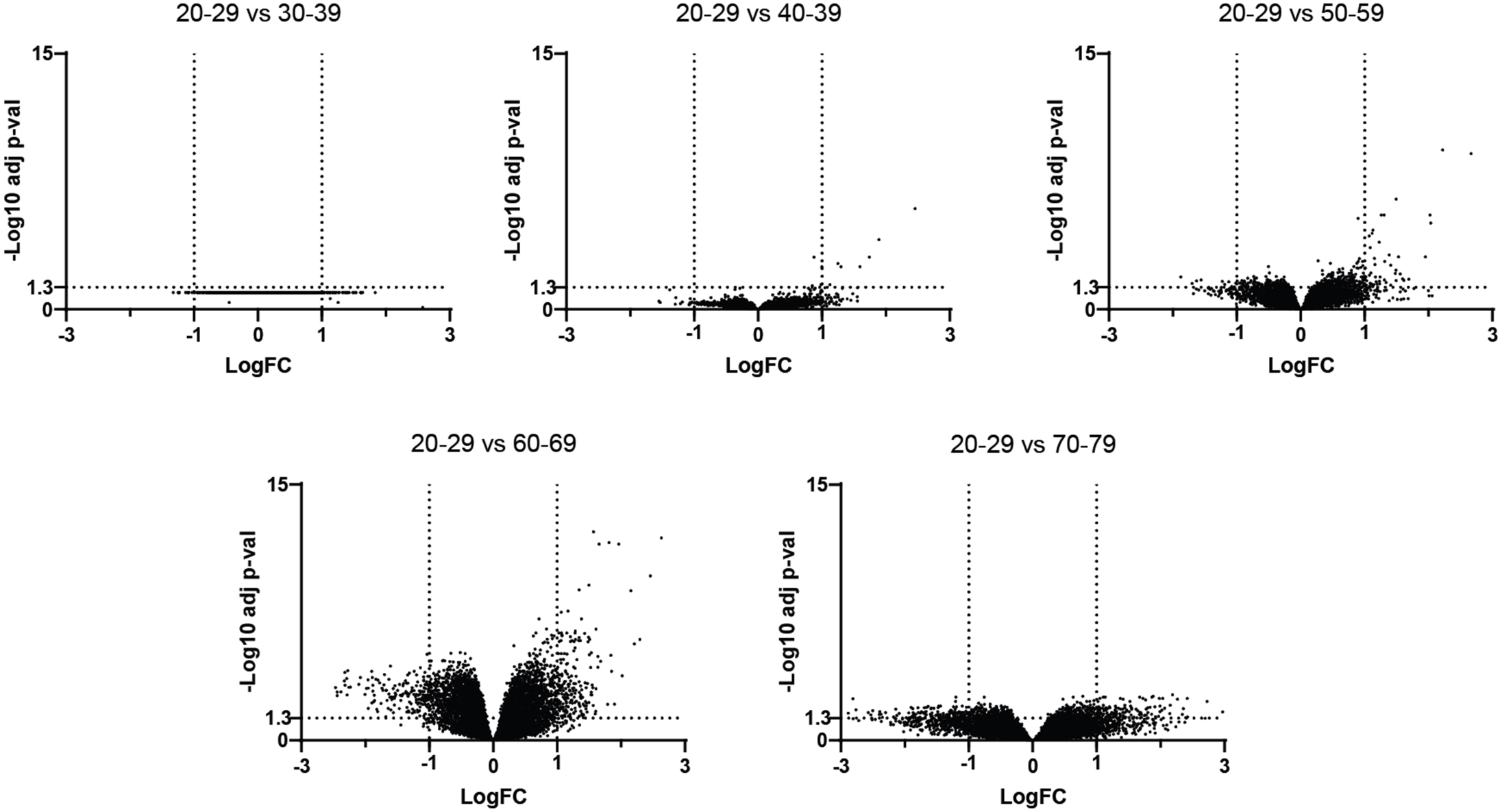
Volcano plot showing lung transcriptional deregulation in 286 GTEx male samples during aging, represented as −log10 (adjusted p-value) and log fold change difference. Dashed lines represent our cutoffs (LogFC ≥ |1| and FDR<0.05). The differentially expressed genes (DEG) with age were identified by distributing the expression data of each individual into five age groups (30-39; 40-49; 50-59; 60-69 and 70-79 years old) and compared with a group of young individuals (20-29 years old). Number of men: 20-29 yo (n = 16), 30-39 yo (n = 21), 40-49 yo (n = 47), 50-59 yo (n = 109), 60-69 yo (n = 86), 70-79 yo (n = 7). Number of women: 20-29 yo (n = 11), 30-39 yo (n = 9), 40-49 yo (n = 29), 50-59 yo (n = 36), 60-69 yo (n = 53), 70-79 yo (n = 3). 7 genes were up regulated on 40-49 yo, 56 on 50-59 yo, 217 on 60-69 yo, and 144 on 70-79 yo. 11 genes were down regulated on 50-59 yo, 179 on 60-69 yo, and 291 on 70-79.

**Figure S2.**
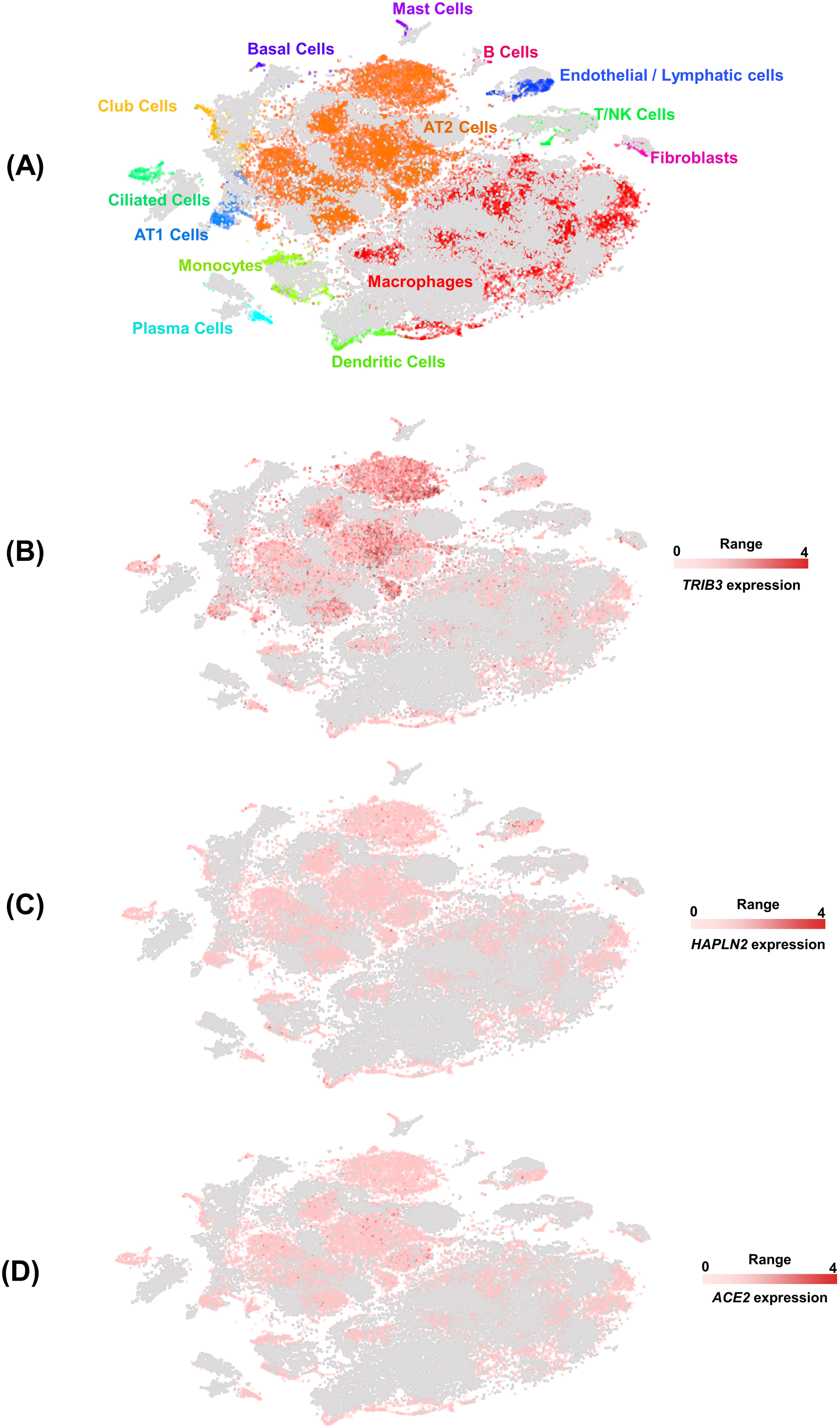
Single-cell gene expression analyses of *TRIB3, HAPLN2*, and *ACE2* in lung cells. A) Unsupervised clustering demonstrates different cell populations identified in non-diseased lung human samples in a *t*-distributed Stochastic Neighbor Embedding (tSNE) plot, as described previously ^20^. Grey dots represent single-cells from pulmonary fibrosis samples that were not included in the present analysis. Single-cell gene expression of *TRIB3* (B), *HAPLN2* (C), and *ACE2* (D) in different cell populations of the lung. The images were generated using the dataset ^20^, available at <u>nupulmonary.org/resources/</u>. The range represents the minimum and maximum expression.

**Figure S3.**
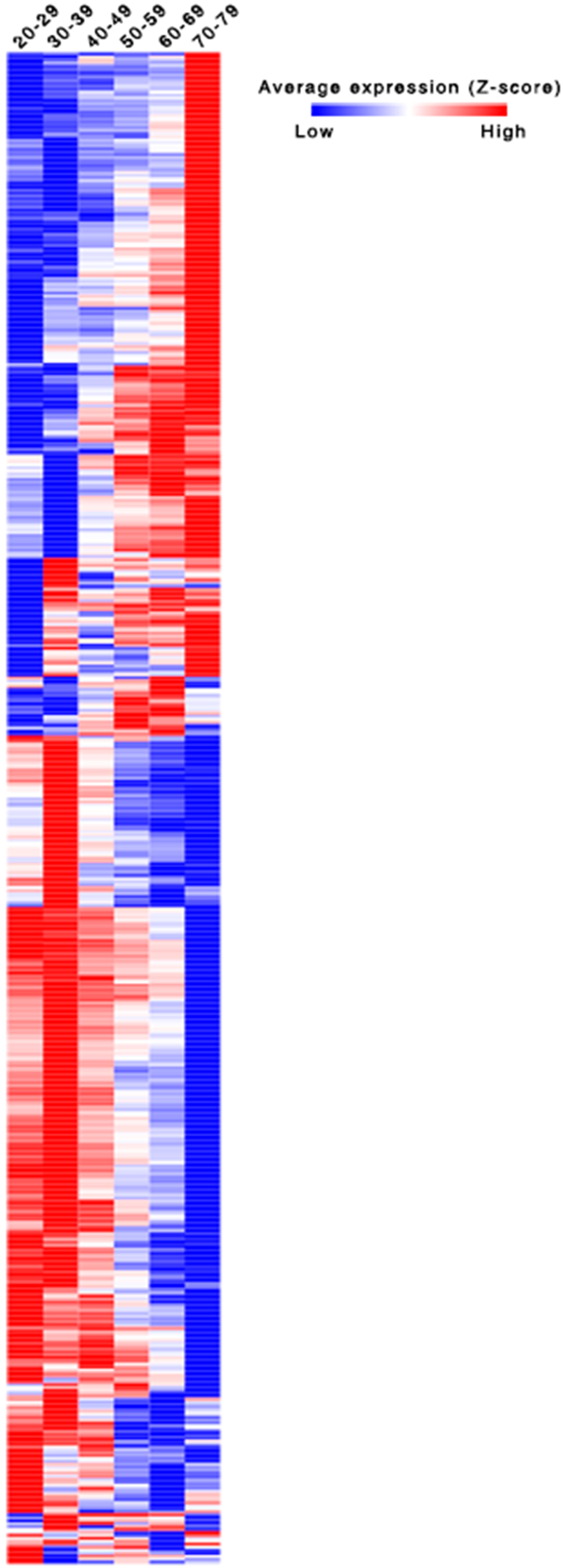
Heatmap with the mean expression of DEGs on female samples (n=129), normalized by the Trimmed Mean of M-values (TMM) and Z-scored by row. Fewer genes clustered on gradients compared with males (Figure 1), which could mean a more substantial noise on the data due to decreased sample size or female hormonal cycle.

